# Behavioural, but not motivational, trade-offs in a cockroach

**DOI:** 10.1101/2025.10.31.685749

**Authors:** Joe Jenkinson, Joanna Logan, Christopher Robertson, Keith Lockhart, David N. Fisher

## Abstract

The ability to flexibly trade off access between competing or antagonistic stimuli (“motivational trade-offs”) is a key criterion for assessing whether animals can feel pain. However, in some commonly farmed, exterminated, and studied insect orders such as Blattodea (cockroaches and termites), the ability to make motivational trade-offs is unconfirmed, which makes it difficult to assess their welfare needs. Here we gave cockroaches (*Blaptica dubia*) a choice between access to shelter or to nutrition across a series of linked experiments to assess whether they are capable of motivational trade-offs and if they are flexibly adjusted due to external conditions and/or injury and develop across ontogeny. We found cockroaches avoided a food reward when it was exposed to light (a behavioural trade-off), but they did not adjust their preference flexibly in response to lighting intensity or increased hunger level (no motivational trade-off). We did find that injury reduces willingness to accept light exposure when accessing nutrition, which is consistent with the idea of the sensation of pain reducing risk-taking behaviour. Finally, we showed that juvenile cockroaches match adults in not making a motivational trade-off. All together our results provide much-needed evidence for criteria of whether insects feel pain and suggest upper limits to cognitive complexity and flexibility in Blattodea. These findings can be integrated with other neurological and behavioural measures to help us judge how we treat insects and other invertebrates in our care.

## Introduction

Animal welfare has varied definitions but typically refers to the quality of life experienced by the animal. Generally, using animals ethically means that negative physical and psychological states should be minimised and positive ones promoted, therefore affording animals with good welfare and a “life worth living” (Hewson 2003; Reimert *et al*. 2023). Considerations include suitable nutrition and physical environment, good health, positive behavioural interactions, and the animals mental state (which is influenced by the factors mentioned beforehand; Mellor *et al*. 2020). Understanding and improving welfare is relevant not only in agricultural settings, but also in scientific research and for wild animals. Researchers have a duty of care to minimise animal suffering when using animals for experimentation or other practical purposes (Russell and Burch 1959), and in wild animals, as welfare can be an important component of conservation (Paquet and Darimont 2010).

In order to be protected under animal welfare legislations there must be sufficient evidence supporting the plausibility of sentience and therefore the capacity to suffer. Key in concluding whether an animal can suffer is whether it feels pain: an unpleasant sensory and emotional experience associated with, or resembling that associated with, actual or potential tissue damage (Raja *et al*. 2020). There is a broad consensus that pain is plausible in vertebrates including birds, amphibians, reptiles, fish, and mammals (Gentle *et al*. 1991; Machin 1999; Sneddon 2003; Sneddon *et al*. 2014; Williams 2019), supported by behavioural indicators such as protective responses, changes in motivation, and avoidance learning following injury (Rutherford 2002). In addition, these pain-feeling animals possess physiological indicators for the ability to feel pain such as nociceptors, complex neurological pathways linking sensory brain regions, and modulation of these nociceptive pathways when exposed to analgesics (Crump *et al*. 2023; Gibbons *et al*. 2022b; Key *et al*. 2021). However, invertebrates (approx. 97% of animal species) such as insects have traditionally been considered to lack key physiological indicators and not to show the behavioural indicators that indicate they feel pain (Adamo 2016, 2019; Eisemann *et al*. 1984), and so most animal welfare legislation excludes invertebrates (but see: Barrett and Fischer 2025). For example, in the UK the only invertebrates protected under legislation during scientific research are cephalopods (*Animals (Scientific Procedures) Act 1986* 2021), which reflects the evidence supporting the possibility they feel pain (e.g.: Crook 2021).

One theory for why feeling pain might be adaptive is that it helps an animal avoid potentially damaging situations in the future (Bateson 1991; Crook *et al*. 2014). The supposed adaptive value of pain suggests that many taxonomic groups may possess pain-sensing mechanisms, not solely vertebrates. Recently, the evidence for behavioural and physiological indicators of the ability to feel pain has been re-assessed for decapod crustaceans and cephalopod molluscs (Birch *et al*. 2021) and for insects (Gibbons *et al*. 2022a). While these frameworks have been criticised for implying all criteria are equally important and lacking a framework for integrating negative evidence into considerations (Diggles *et al*. 2024), they provide a useful roadmap for assessing different aspects of pain sensation. What Birch et al. (2021) and Gibbons, Crump et al. (2022) report is that, when scientists have looked for the presence of the physiological and behavioural indicators in these taxa (often they have not), they find them, making it much more plausible that insects may feel pain. However, gaps in evidence remain, limiting our ability to be decisive about whether invertebrates are likely to have the capacity to suffer and if so, what legal protections are required for their welfare.

The behavioural indicators of pain are each more than a mere reflex and demonstrate some longer-term shift in response to a noxious stimulus (Bateson 1991; Sneddon *et al*. 2014). For example, motivational trade-offs require an animal to demonstrate a flexible trade-off between competing stimuli, such as accepting exposure to a noxious stimulus to gain a reward or avoid an even more noxious alternative (Brown and Birch 2025). Ideally, it should be clear that the trade-off is based on a mental representation, such as a memory of a stimuli, rather than direct experience (Gibbons *et al*. 2022c). Motivational trade-offs are thought to stem from adaptive decision making, where animals in the wild must weigh up several competing factors against each other e.g., shelter availability vs predation risk (Lima and Dill 1990). The presence of motivational trade-offs across different domains, showing the processing of different types of information into a common currency, is suggestive of central processing of stimuli (Crump et al., 2022, although further follow up evidence from neurological tests should be sought to confirm this), which is important for inferring more complex cognition. One example of a motivational trade-off in invertebrates comes from *Drosophila melanogaster*, which show some willingness to tolerate 120V shocks to gain a reward of ethanol but not to gain the less-rewarding sucrose (Kaun *et al*. 2011). However, in insects motivational trade-off research has focused on *Drosophila melanogaster*, *Apis mellifera*, and *Bombus terrestris*; these three species may not represent the millions of other insect species, leaving a knowledge gap across other species and orders (Gibbons *et al*. 2022a).

One well-studied insect order without evidence for motivational trade-offs is Blattodea (cockroaches and termites). Blattodea are exceptionally widespread, featuring in nearly all climates with several species being well-known as domiciliary pests and disease vectors (Bell *et al*. 2007), and several cockroach species being increasingly used in production settings (Siddiqui *et al*. 2024). Cockroaches are also widely used in neuroscience research (Huber *et al*. 1990). Despite these factors, there are no studies testing whether cockroaches make motivational trade-offs (although Varnon and Adams (2021) did find that the presence of food inhibited startle responses to light in the orange head cockroaches (*Eublaberus posticus*), suggesting the ability to integrate information across domains if not evidence for a trade-off between accessing competing goals). We aim to fill the gap around motivational trade-offs in Blattodea here. Additionally, we explored motivational trade-offs in cockroach nymphs, as Gibbons *et al*. (2022a) highlighted that no juvenile insects have been tested for the ability to make motivational trade-offs. It is crucial that juveniles are tested, as many insects (such as yellow mealworm beetle [*Tenebrio molitor]* and black soldier fly [*Hermetia illucens*]) spend the majority of their lives, especially in production settings, as juveniles, and so their capacity to suffer in this life stage is key.

We carried out three linked experiments to assess the extent of motivational trade-offs in adult and juvenile cockroaches. We used a trade-off between access to nutrition (a positive stimulus) vs. exposure to light (a negative stimulus) to test for motivational trade-offs. First, we tested whether conditions altered the magnitude of the trade-off between access to nutrition and exposure to light. Demonstrating the flexibility of motivational trade-offs is key for understanding whether these are simple reflexes or evidence of central processing. Second, we determined if an injury altered this trade-off. Injury altering subsequent behaviours that indicate an increase in protective behaviour is considered key for showing the feeling of pain rather than simply nociception (Sneddon *et al*. 2014). Finally, we determined if juveniles made the trade-off, and if the magnitude of the trade-off varied with mass, which we used as a proxy of developmental stage (Wu 2013). We used the Blaberid cockroach *Blaptica dubia*, a species widely used in the pet trade as live feed but also increasingly used in scientific research. We predicted that cockroaches would trade-of between access to nutrition and exposure to light, and that the magnitude of the trade-off would vary in response to conditions and be influenced by injury, indicating a motivational trade-off rather than merely a behavioural trade-off. For juveniles, we predicted they would make motivational trade-offs and further predicted that larger individuals would be more likely to perform trade-offs as the need for resources with larger moults is more pronounced than when smaller and as their cognitive complexity is expected to increase as they develop. While no one experiment is conclusive evidence of the ability to make motivational trade-offs, we aim to provide three lines of evidence, allowing us to arrive at a general conclusion on the probability that cockroaches are capable of motivational trade-offs (Brown and Birch 2025).

## Methods

### Stock and housing

We used a population of *B. dubia* that have been maintained at the University of Aberdeen since 2021, with periodic purchase of new individuals to limit inbreeding depression. The *B. dubia* stock population are stored in an insectary room within 48L plastic (polypropylene) source boxes (60cmx40cmx35cm, Really Useful Boxes, Really Useful Products Ltd.) each containing several 24-egg sized cardboard carton for shelter, which are replaced after signs of degradation. Source boxes are cleaned twice weekly with 70% ethanol solution. Hydration in the form of three 40g carrot slices and food in the form of 10g of Sainsbury’s Complete Nutrition Adult Small Dog Dry Dog Food (rough nutritional values: 1527kJ energy, 24g protein, 12g fat per 100g) are also replaced twice per week. This insectary remained between 28-30°C under 50% humidity with a 12:12 light/dark light system that kept lights on only between 7am and 7pm (further details on husbandry are provided in Fisher 2023).

### Base Method

We conducted three different experiments using the same base method, with differing variables explored in each experiment to test for the existence of motivational trade-offs and their extent in *B. dubia*. In each experiment we tested for motivational trade-offs by comparing the willingness of individuals to stay in one side of a box that was empty, but sheltered, or go to the other side of the box that contained a piece of carrot, which was one of two treatments: either exposed to light (“exposed”) or sheltered (“sheltered”). By comparing these two groups we can determine if how much the cockroach goes to the carrot depends on being exposed to or sheltered from light, indicating a behavioural trade-off between access to shelter and access to nutrition. Further, we can vary the experimental conditions or characteristics of individuals to determine the flexibility of the trade-off. It is testing the interaction between changing experimental conditions and exposed or sheltered that is key for testing for the presence of motivational trade-offs, as it demonstrates the integration of information from different sources (Read and Nityananda 2026). Experiment 1 explored the impact of conditions (light intensity or hunger), experiment 2 the impact of injury, and experiment 3 the impact of development. We only used males for our experiments with the adults as females from the stock population were being used in other experiments and it was easier to injure the males for experiment 2 due to their exposed wings. We did not sex the nymphs for the experiment on juveniles and so assume the experiment included an equal number of males and females. We filmed all trials with ABUS IP video surveillance 8MPx mini tube cameras (ABUS, Germany) through the “AnyCam” software (AnyCam.iO) rather than carry out direct observations, to limit the impact of observer presence on behaviour.

In the first step of the base method, we randomly selected cockroaches from their 48L source boxes and placed them into a separate environment without access to nutrition or hydration for 24 hours. We did this to induce thirst and/or hunger in the cockroaches, increasing their motivation to leave the always-sheltered side of the box and access the carrot. For experiment 1, we placed cockroaches separately into small boxes (7.9cm x 4.7cm x 2.2cm), while for experiments 2 and 3 individuals were held together in a 48L box. After the 24 hours we transferred each individual cockroach to a 0.9L box (22cm × 10cm x 7cm) that was either 50% or 100% shaded depending on whether they were in the exposed or sheltered treatment respectively (Fig. 1). Shade was created using plain white A4 paper that was cut to shape and taped around the top and sides of the boxes excluding one side to allow lateral visibility into each box. In each box we placed a piece of carrot of approximately 3g. In experiment 1 we added carrots to the boxes before the cockroaches were taken to the recording room, whereas for experiments 2 and 3 we added their carrot slices at the point filming began. These 0.9L boxes were cleaned with 70% ethanol between trials to limit any effect of scent or contaminants on the next cockroach to enter the box.

**Figure 1.**
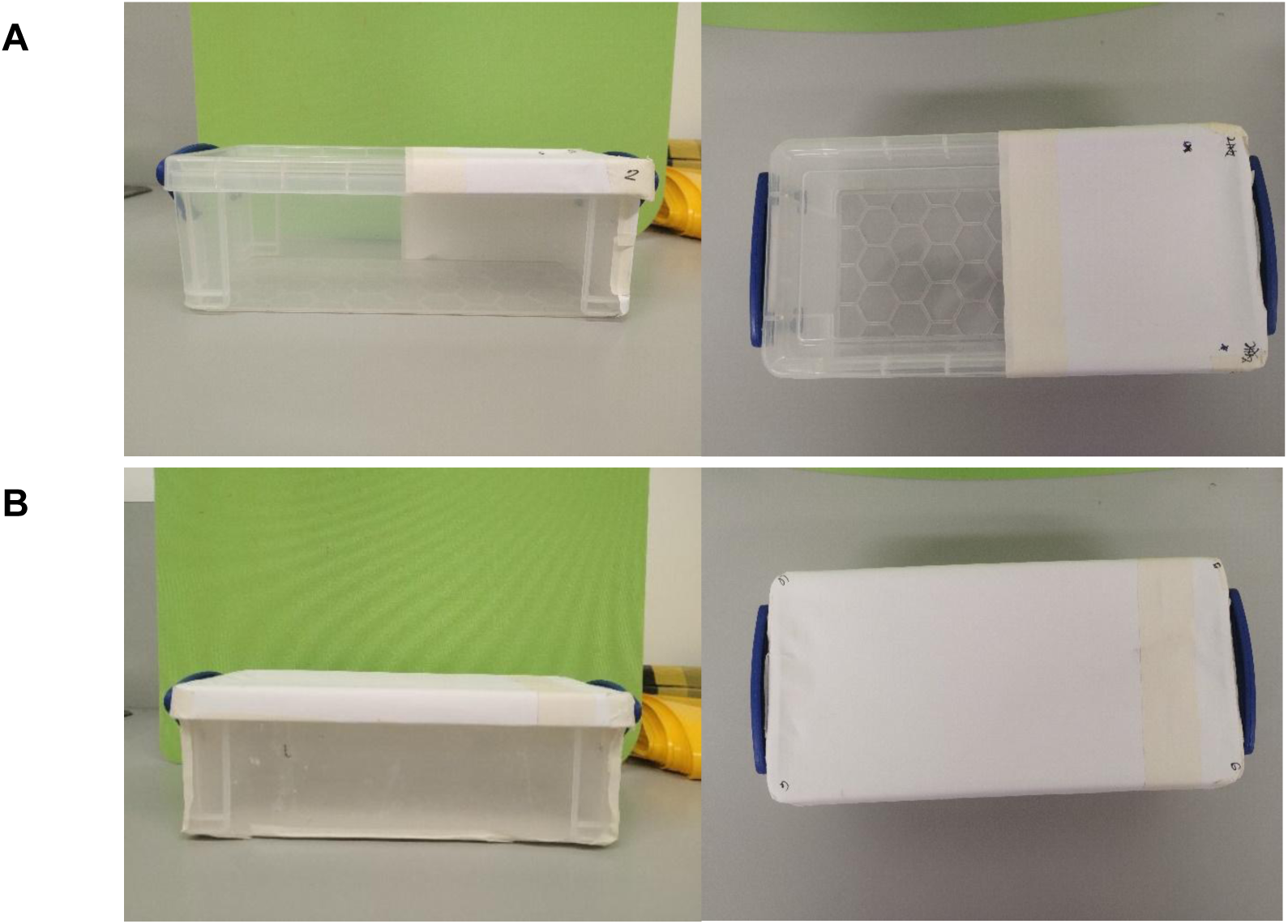
Lateral (left) and dorsal (right) views of the (A) Exposed box and (B) Sheltered box (filming is conducted from top-down).

We transferred cockroaches in individual 0.9L boxes to the recording room, with an equal mix of exposed and sheltered treatments. We heated the recording room with portable heaters to approximately 25°C for at least an hour before beginning any trials, while we recorded temperature during trials with a thermometer and controlled for its effect statistically (see Data analysis). As the recording room was not sealed, to prevent cockroach escape into the wider building we placed each 0.9L trial box into a larger box for filming. For trial 1, we placed two randomly ordered pairs of 0.9L boxes in larger 48L boxes. We placed a single camera per 48L box (covering two 0.9L boxes) 6.6 cm above the ground at a distance of 30cm from the 0.9L boxes (Fig. 2A&B) and used up to four cameras simultaneously to assess up to eight individuals at once. For trials 2 and 3 we placed the 0.9L boxes singly into 5L boxes for recording, using one camera (elevated by 14cm) per four boxes by filming two side-by-side and stacking an extra pair of empty 5L boxes behind the first two to prop up the next two (Figure 2C&D). We used up to two cameras simultaneously allowing us to assess up to eight individuals at once. We positioned these cameras to provide a clear side view of all the boxes in front so that all cockroaches and their position were always clearly visible in recordings, and since the conditions for the cockroaches do not change between the set-ups (they all experience a 0.9L box with a piece of carrot and either exposed or sheltered treatment) we believe the experiments to be comparable. We randomly assigned cockroaches to the camera and location in the arrangement we placed them in. We placed a pet heat mat (models TK-HPP7040A & TK-HPP6540, OnKey Electronic Technology Co. Ltd) underneath each of the paired 5L or individual 48L boxes containing cockroaches (Fig. 2) to maintain them at a recommended temperature of ∼30^°^C so that low temperatures did not influence their locomotive ability (Alamer and Hoffmann 2014).

**Figure 2.**
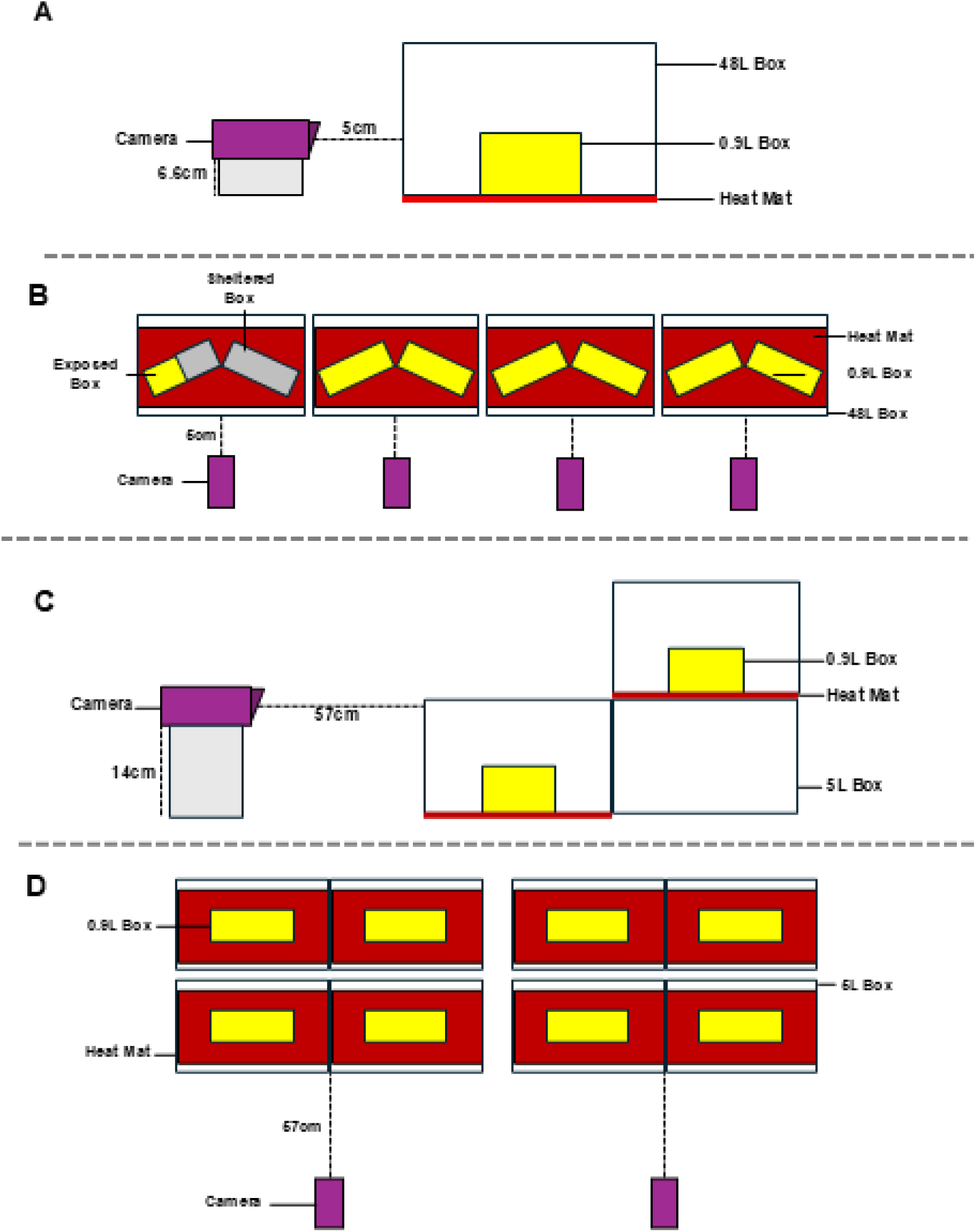
Diagram of experiment set up for experiment 1 lateral (A) and dorsal (B) views and experiments 2 and 3 lateral (C) and dorsal (D) views. Each set up was designed to give a clear view of the open side of each 0.9L box so that the position of the cockroach within could be determined.

**Figure 3.**
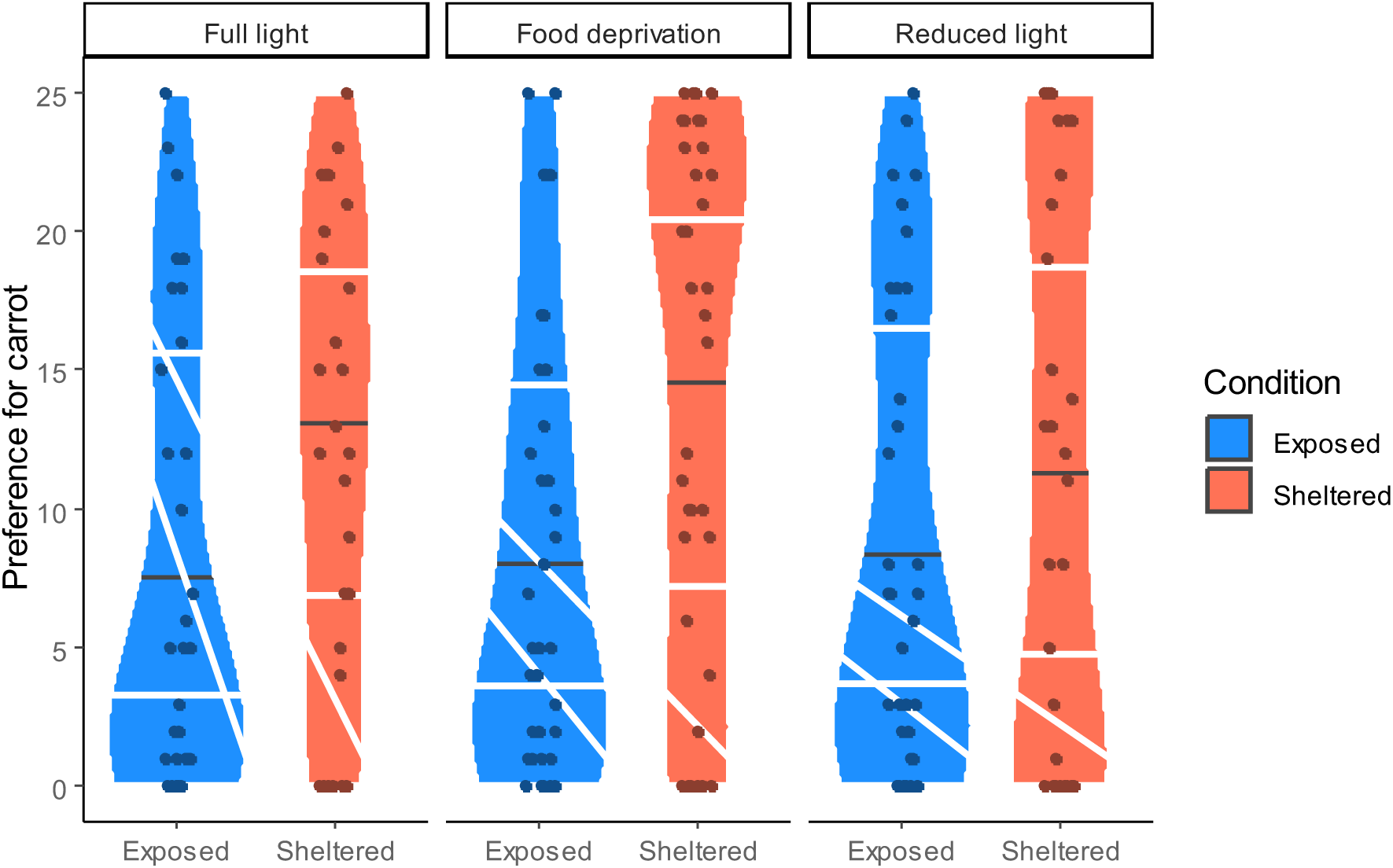
Dot and violin plots showing the cockroaches’ relative preference for carrot (number of times with carrot [max 25]) in exposed (blue) and sheltered (orange) arenas from experiment 1. We show the results for each experiment separately to illustrate how the differences between exposed and sheltered were consistent across the experiments (full light, 48-hour food deprivation, and reduced light from left to right). Black lines show the medians, white lines the 25^th^ and 75^th^ percentiles. Width of the “violins” indicates the relative density of points.

We set the cameras to record for 30 minutes and then immediately left the recording room. At the end of experiment 1 we transferred cockroaches back to the source box they came from. Cockroaches could thus have been used more than once between different trial days but would not be used in the same trial day or consecutive days due to being preselected for isolation the day prior. This provided time for readjustment to their source box environments after trials before potentially being used again on another trial day, negating any behavioural impacts. In experiment 2 all cockroaches were frozen in −20°C after the experiment, which is an effective means of euthanasia in this species (100% mortality after one hour; Tucker *et al*. 2023). Approaches using anaesthesia were not available to us at the time. The use of euthanasia means that individuals in this experiment were never re-used. In experiment 3 nymphs were returned to the stock population but placed into a specific box, and so were not re-used in the experiment.

### Experimental Conditions

In experiment 1 we ran three separate trials, all underpinned with the base method. The first trial was only the base method. In the second trial we then tested the impact of conditions by altering the light level in the open half of the exposed boxes by attaching translucent paper to partially block light over the food as opposed to allowing full light. Third, we explored the effect of changing the starvation period from 24 hours to 48 hours. The experimentation period (including laboratory testing, video extraction, and filming) ran from 11/10/2022 to 22/11/2022 and used 72 individuals, although recordings were lost for 11 of these, leaving 61.

For experiment 2 we used the base method, but after the 24-hour starvation period all the starved cockroaches were randomly allocated as injured or uninjured. For the injured individuals, we cut off 1cm of their outer wings with sharp scissors. The wings were clipped as *B. dubia* wings in males have limited function and so wouldn’t impact locomotion for the experiment, but contain nerves and so may be able to sense damage (Kniazeva 1976; Pipa and Cook 1959). Recently we have shown male *B. dubia* with this type of injury perform a range of behaviours directed at the site of the injury that uninjured individuals do not, strongly suggesting the damaged is sensed (Farquhar and Fisher 2026). We handled uninjured cockroaches for around 40 seconds as a “sham” treatment, which is the approximate time the injured cockroaches spent being handled while having their wings clipped. Experimentation ran from 21/10/2024 to 12/11/24 and used 120 individuals, but data from 16 trials were lost due to camera failure, leaving 104.

For experiment 3 we used body mass as a proxy for age/developmental stage as instars differ substantially in mass (Wu *et al*. 2017). We weighed each nymph to be used in the experiment before the starvation period using a Fisherbrand Analytical Balances, readability 0.0001g scales. Body mass was recorded to two decimal places. Experimentation ran from 28/10/2024 to 20/11/24 and used 80 individuals.

### Video and data analysis

For each experiment we followed a similar analysis approach. We first removed the first 5 minutes of each trial as during this “habituation” period the cockroach may not have been responding to the experimental treatments but rather to the novelty of the box. We watched the remaining video and every 60 seconds (experiment 1) or every 30 seconds (experiments 2 and 3) counted the number of times during the trial the cockroach was either on the same side of the box as the carrot, the opposite side, in the middle, its side could not be determined (experiments 2 and 3 only), and the number of times it could not be seen (experiment 3 only). We increased the sampling rate for experiments 2 and 3 to give a higher resolution on behaviour.

For experiments 1 and 2 there were no time points where the cockroach could not be seen, and so both of these were analysed in the same way. For experiment 1 analysed data from all three conditions (base method, reduced light, longer starvation) together. Initial analyses revealed both overdispersion and a high frequency of zeros. We therefore used the package “glmmTMB” to fit a GLM with a negative binomial error family with a linear parametrisation (Hardin and Hilbe 2007) and a zero-inflation term. We fitted the number of time points the cockroach was on the same side as the carrot as the response variable, and predictor variables of exposed/sheltered as a two-level factor, conditions (full light/reduced light/starvation) as a three-level factor, the interaction between these two terms, and temperature (scaled to a mean of 0 and a variance of 1, which aids the fitting and interpretation of regression models; Schielzeth 2010) as a continuous covariate. For experiment 2 we again fitted a negative binomial GLM, a zero-inflation term, and the same response variable, with predictor variables of exposed/sheltered as a two-level factor, injured/uninjured as a two-level factor, the interaction between these two terms, and temperature (again scaled to a mean of 0 and a variance of 1) as a continuous covariate.

For experiment 3 there were eight trials where the cockroach could not be seen at four time points, and so we wanted to account for a different number of overall observations. We therefore initially fitted a binomial GLM with number of times the cockroach was the same side as the carrot as the “successes” and the total number of observations where the individual could be seen as the number of “trials”, which accounts for variation among cockroaches in the latter value. However, this model was always overdispersed across a range of error distributions. Instead, we calculated the proportion of times the cockroach was on the left-hand side (where the carrot was), and entered that as a response variable into Ordered beta regression GLM (Kubinec 2023). As this fits continuous proportion data in the interval 0, 1, we did not need to include a zero-inflation term (the model had satisfactory dispersion with and without the zero-inflation term, so we used the simpler model without). We included predictor variables of exposed/sheltered as a two-level factor, mass in grams as a continuous variable (scaled to a mean of 0 and a variance of 1), the interaction between these two terms, and temperature (again scaled to a mean of 0 and a variance of 1) as a continuous covariate.

For all models the interaction between exposed/sheltered and the other predictor of interest was not significant, and so we calculated p values with a type II sum of squares ANOVA in the package car (Fox 2016). The main effect of exposed/sheltered indicates whether the cockroach is more or less willing to go to the carrot when it would be exposed to light in doing so, providing evidence of a behavioural trade-off. Meanwhile the interaction between exposed/sheltered and the other factor in each analysis would have indicated whether this trade-off depended on conditions (experiment 1) or injury (experiment 2), and so had the flexibility required for a motivational trade off, while the interaction with mass indicated whether this trade-off changed with age (experiment 3).

### Ethical note

While no formal ethical approval was required for this work, we adhered to ASAB guidelines throughout. Our approach was minimally invasive aside from in experiment 2 where we were required to injure the cockroaches to test our hypothesis. These individuals were tested immediately after injury and euthanised directly after the experiment to minimise any period of potential suffering. We also deprived the cockroaches of food and hydration for 24 or 48 hours, but this is well within conditions cockroaches are naturally exposed to and are specifically adapted for (Bell *et al*. 2007). At the end of each study animals were returned to the stock population unless stated otherwise.

## Results

### Experiment 1 (impact of conditions)

Cockroaches avoided the carrot more when it was exposed to light (main effect of exposed/sheltered, incidence rate ratio [IRR] = 0.519, χ^2^_1_ = 31.704, p < 0.001), indicating a behavioural trade-off. The magnitude of this trade-off did not vary among conditions (interaction between exposed/sheltered and condition, χ^2^_2_ = 0.604, p = 0.739) and the average preference for carrot did not differ among conditions (main effect of conditions, χ^2^_2_ = 0.788, p = 0.674). Cockroaches’ preference for carrot was unaffected by temperature (β = −0.064, χ^2^_1_ = 1.437, p = 0.231). The dispersion parameter was 5.04 and the zero-inflation term had an estimate of −1.324 with a standard error (SE) of 0.181.

### Experiment 2 (impact of injury)

In experiment 2, cockroaches again showed a behavioural trade-off, avoiding the food more if it was exposed to light (main effect of exposed/sheltered, IRR = 0.751, χ^2^_1_ = 11.412, p = 0.001). Additionally, injury reduced their willingness to go to the carrot (main effect of injury, IRR = 0.787, χ^2^_1_ = 8.817, p = 0.003). However, the trade-off due to light was unaffected by injury (interaction between exposed/sheltered and injury, χ^2^_1_ = 0.941, p = 0.332; Fig. 4). Cockroach preference for carrot was unaffected by temperature (β = 0.019, χ^2^_1_ = 0.110, p = 0.741). The dispersion parameter was 6.76 and the estimate for the zero-inflation term was −4.648 with an SE of 1.018.

**Figure 4.**
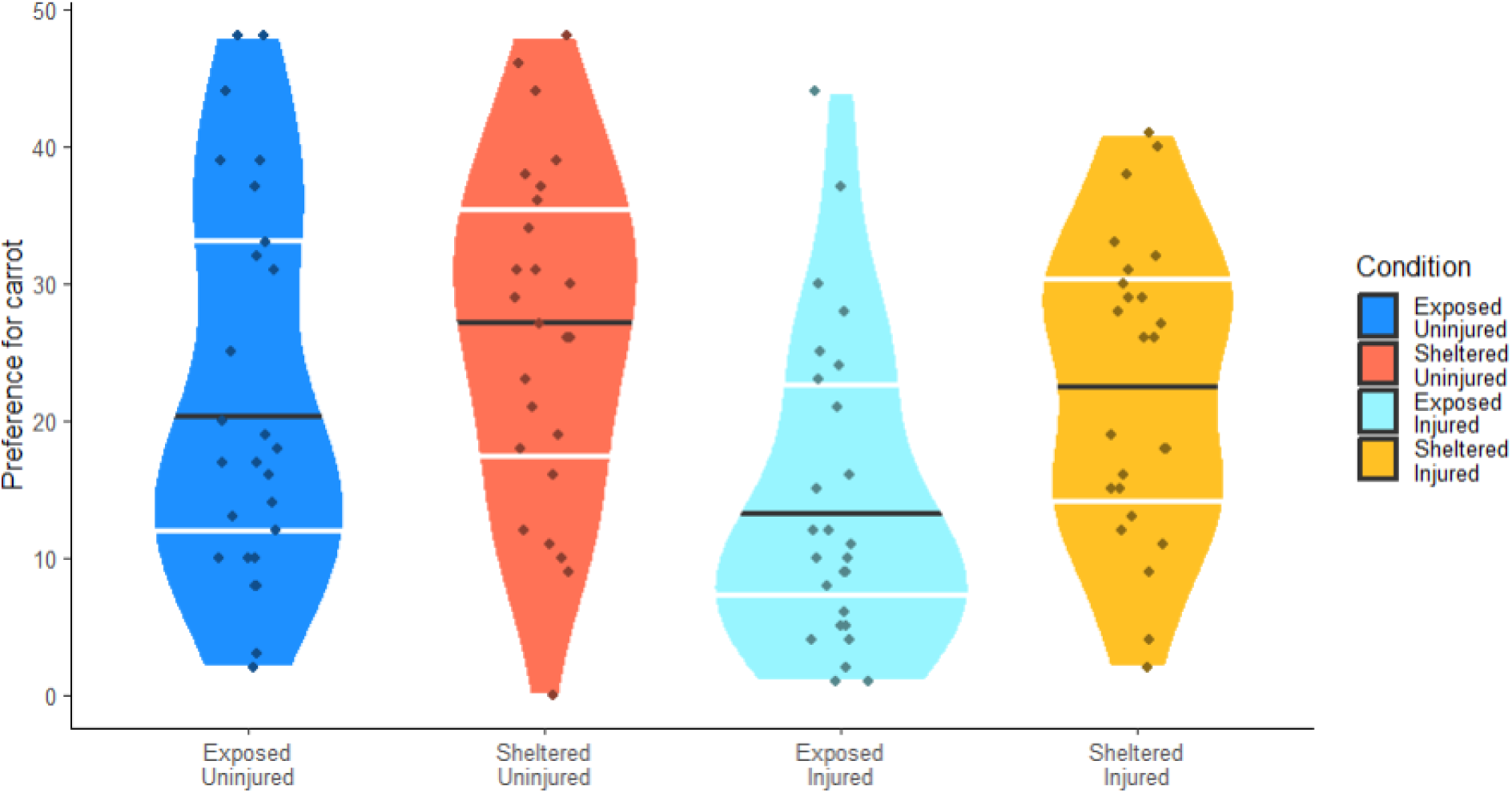
Dot and violin plots showing the cockroaches’ relative preference for carrot (number of times with carrot [max 50]) in exposed or sheltered arenas and when injured or uninjured from experiment 2. Dark blue shows exposed and uninjured, orange sheltered and uninjured, light blue exposed and injured, yellow sheltered and injured. Black lines show the medians, white lines the 25^th^ and 75^th^ percentiles. Width of the “violins” indicates the relative density of points.

### Experiment 3 (impact of development)

Experiment 3 showed nymph cockroaches also make a behavioural trade-off and avoid light (main effect of exposed/sheltered, log-odds ratio = 0.420, χ^2^_1_ = 6.952, p = 0.008). Heavier individuals showed reduced preference for carrot (β = −0.180, χ^2^_1_ = 4.087, p = 0.043). The behavioural trade-off did not change with mass (interaction between exposed/sheltered and mass, χ^2^_1_ = 0.575, p = 0.448; Fig. 5). Cockroach preference was again unaffected by temperature (β = −0.120, χ^2^_1_ = 0.639, p = 0.424). The dispersion parameter was 1.39.

**Figure 5.**
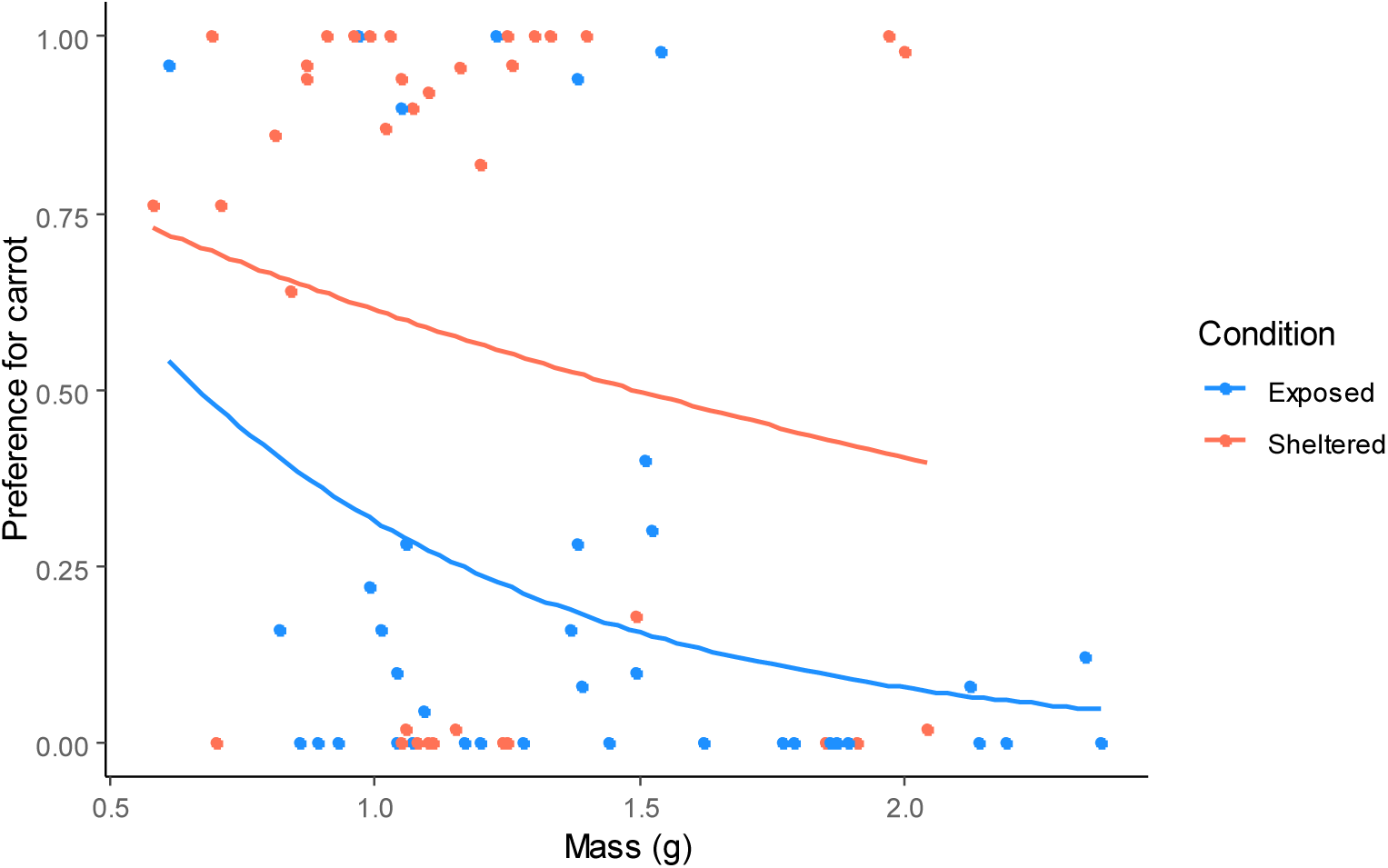
Scatterplot showing the cockroaches’ relative preference (number of time points with carrot/total time points observable) for carrot in exposed (blue) and sheltered (orange) arenas across masses (developmental stages) from experiment 3. The difference between exposed/sheltered did not clearly change across masses. Trend lines were drawn in ggplot2 (Wickham 2009) using a Poisson GLM (with the proportion as a response variable, which differs from our analysis in the main text using an ordered beta regression model as this error distribution is not available in ggplot2, but should still give an illistrative trend line).

## Discussion

We found comprehensive evidence that *B. dubia* cockroaches can make behavioural trade-offs, avoiding food when it was exposed to light. However, the strength of the trade-off was unaffected by the strength of the light or the relative value of the nutrition, indicating no motivational trade-off. When we injured cockroaches, they were less willing to go to the nutrition, and this effect was additive with the light level, indicating some response to injury but no integration into other decision-making processes. Finally, the trade-off between nutrition and shelter was present in nymphs but did not change with mass, suggesting no development/change across ontogeny.

Animals that react to negative stimuli through more than reflexes but with flexible behavioural adjustments that mitigate their risk of future threats are thought to be more likely to have the capacity to feel pain and therefore suffer (Gibbons *et al*. 2022b; Sneddon *et al*. 2014). Simple behavioural trade-offs however can be produced by cognitively simple animals and do not imply the integration of diverse sources of information into a common currency (Read and Nityananda 2026). The lack of flexibility in the trade-off between exposure to light and access to nutrition we observed here therefore provides no evidence for the ability of Blattodea of make more complex motivational trade-offs, as the relative value of the food reward (manipulated by a longer period without food) or the relative value of the light risk (manipulated by altering light levels) did not alter the cockroaches’ behaviour. While there is evidence that Blattodea meet six out of eight criteria for feeling pain (Gibbons *et al*. 2022a), our study does not support more complex decision making around motivations. The presence of trade-offs alone are not evidence for feeling pain (Brown and Birch 2025), and similarly negative evidence for this criterion does not eliminate the possibility that cockroaches feel pain, but it does encourage us to develop a more nuanced understanding of how different taxa might meet some thresholds but not others, and so how our treatment of them should be sensitive to this.

Our second experiment found injured cockroaches were less willing to access nutrition in general. However, the trade-off between access to nutrition and exposure to light was not more or less pronounced in injured cockroaches, suggesting that damaged cockroaches do not regard the environment as especially risky or themselves as especially vulnerable; what we would expect if feeling pain was adaptive (Elwood 2011). Note that since we injured cockroaches on their wings, which they do not use for locomotion, we can be confident that they did not access the nutrition less simply because their movement was impaired. Past work on American cockroaches (*Periplaneta americana*) has shown they will attend to and groom the site of an injury (e.g., an abdominal puncture wound; Hentshel & Penzlin, 1982) while our study adds to the picture that injury leads to more than nociception because we see an impact on behaviour that lasts well beyond the damaging event (see also: Farquhar & Fisher, 2026, who test the final unexplored criterion of the Birch framework for feeling pain: preference for analgesics when injured). Our finding that wing damage impairs feeding also has implications for insect housing conditions – keeping certain species at high densities often leads to wing damage (Wehmann *et al*. 2022), which we have shown here could lead to reduced food intake – possibly a problem in production settings where high feed intake is key for growth. Studies comparing housing conditions for other forms of damage besides wings would enable us to identify the best conditions for limiting potentially costly damage.

In our final experiment, we showed that *B. dubia* nymphs also make a trade-off between access to nutrition and exposure to light. We found that the magnitude of the trade-off between access to food and exposure to light did not change with mass (which we used as a proxy for developmental stage). This suggests that the requirements to assess conditions and make a suitable trade-off, such as the ability to sense the light and food, as well as the perception of risk posed by the light, are all already present at the ontogeny stages we tested. Exploring these criteria in juveniles is key as research on pain in juvenile insects is minimal compared to that in adults: Gibbons *et al*. (2022a) identified no studies on motivational trade-offs in juvenile insects, and far less research on all eight criteria in juveniles in general. Given many insects spend most of their life as juveniles, before a short period reproducing as adults, and that in some production settings, the vast majority of individuals never reach adulthood before being slaughtered, testing criteria for pain in pre-adulthood is especially important. Our study is just a first step, further work on juveniles in a wider range of holometabolous species is also required to assess the general capacity for insects with this development mode to make both behavioural and motivational trade-offs. Additionally, research is needed in commonly farmed species such as mealworm (*T. molitor*) and black soldier fly larvae (*H. illucens*) which spend the majority of their lives in captivity as larvae.

In summary, we repeatedly showed that *B. dubia* cockroaches make behavioural trade-offs but found no evidence across three separate experiments that they can make the more complex motivational trade-offs indicative of the capacity to suffer. Our results add to the growing body of work assessing how insects and other invertebrates experience the world and will help us draw up suitable recommendations for their housing and handling. Such recommendations will allow their continued beneficial use while acknowledging our growing understanding of how they experience the world.

### Animal welfare implications

Providing no evidence for one criterion of whether insects feel pain adds depth to the emerging image of their capacity to suffer. While injury affects cockroach behaviour beyond any initial reactions, the lack of integration of different sources of information into a single currency suggests limited central processing, which differs from how we expect painful stimuli to be dealt with. We do recommend that housing conditions are assessed to minimise injury to animals, as otherwise behaviours such as feeding and general locomotion could be reduced. We also urge the development of better and nuanced guidelines and new legislation to reflect evolving scientific knowledge on the capacity of insects and other invertebrates to suffer or not (e.g. Fischer *et al*. 2024) reflecting how they may experience the world in ways both similar and dissimilar to more familiar vertebrates.

## Data availability

Data and R code to re-create the analyses presented here can be accessed on FigShare via: https://figshare.com/s/4f7ad21e2dcf33186389.

